# A record-breaking heatwave reduces breeding success and impairs growth in a wild bird population

**DOI:** 10.64898/2026.08.29.747719

**Authors:** David López-Idiáquez, Devi Satarkar, Ben C. Sheldon

## Abstract

Most evidence of the consequences of climate change in natural systems has focussed on shifts in mean temperature^1,2^, but the effects of extreme climatic events (ECEs) remain far less understood. This is particularly true for very severe ECEs that may occur only once every few decades. Understanding the consequences of these severe events for natural populations is nonetheless critical, since their frequency is predicted to rise under current climate change^3^. Here we combine a unique long-term dataset spanning almost five decades of breeding (>20,000 events) and morphological data (>120,000 observations) in adult and nestling great tits (*Parus major*) and blue tits (*Cyanistes caeruleus*) with fine-scale temperature records to examine the effects of an unprecedented heatwave in May 2026 on breeding success and morphology. Average temperature during the heatwave (22-29 May 2026) was 7.85°C above the historical record, reaching +10.5°C (+4.32 SD) at its peak (25-26 May). These record-breaking temperatures significantly reduced adult breeding success and nestling mass relative to expectation in the absence of a heat-wave. Given the heatwave was widespread (Fig. 1A), our findings from a single, exceptionally well-studied population are likely to generalise to other species exposed to the same event, providing key evidence that severe ECEs can substantially harm wild populations.

---

May and June 2026 brought an unprecedented spell of heat across the UK and western Europe, driven by a persistent high-pressure system, with an estimated 14,000 human heat-related deaths across Europe^3,4^. Wild animal populations faced similarly exceptional conditions, as illustrated by our data from Wytham, near Oxford, home to one of the longest-running bird population studies in the world (Fig. 1A, B). Hourly data from thermo-loggers deployed since 2023 show that between 22 and 29 May, mean temperature in Wytham was 6.89°C higher than the average for the same period in 2023-2025, rising to 9.42°C (+3.16 SD) above average during the peak (25-26 May; Fig. 1B). Comparison with historical data from the Central England Temperature dataset illustrates the extreme nature of the event. During the heatwave, Wytham temperatures were 7.85°C above the 1772-2025 mean for the same dates, reaching +10.5°C (+4.32 SD; Fig. 1B) at the peak, which corresponds to an event expected less than once every 50 years. These record-breaking temperatures in Wytham are consistent with records elsewhere across the UK (Fig. 1A) and continental Europe^4^, underscoring the exceptional and widespread nature of the event.

**Figure 1:**
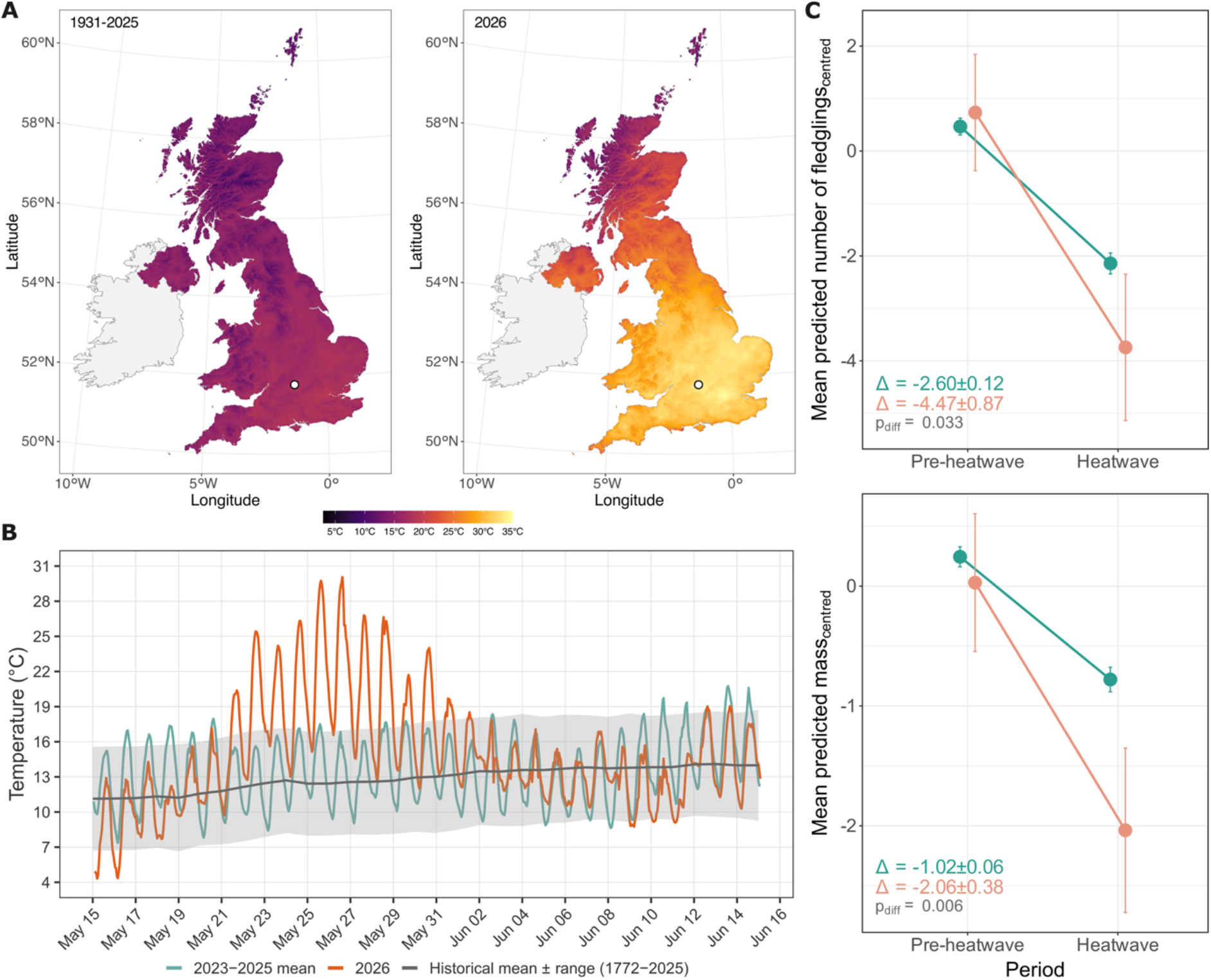
A) Mean daily maximum temperature in the United Kingdom during the 25 and 26 of May between 1931 and 2025, and in 2026. The white dot represents Wytham Woods, near Oxford. Data from HadUK-Grid 1km gridded observational dataset (v.1.3.2). B) Temperature fluctuations in Wytham Woods between 15 May and June 15 for 2023-2026. Orange and green lines represent mean hourly temperatures from 218 temperature loggers in 2026 (orange) and 2023-2025 (green). The grey line and ribbon represent the mean daily temperature ± mean daily temperature range from the Central England Temperature dataset (CET; 1772-2025). C) Heatwave effects on great and blue tit fledgling number and nestling mass. Dots and whiskers represent predicted mean ± standard deviation values from the models. Pre-heatwave and heatwave represent the relative periods before and after the heatwave (see SI-1). Orange represents the heatwave year (2026) and green the non-heatwave years (1978-2025). Values within the plot represent the mean change in the dependent variable between periods and its standard error. The p-value denotes the significance of the difference between heatwave and non-heatwave years. Number of fledglings and nestling mass were centred to a mean of zero.

The heatwave coincided with the breeding season for many birds, a period of peak energetic demand when parents may be particularly vulnerable to environmental extremes. ECEs in this and other systems have typically been defined as days with mean temperatures 4.5-5°C above the monthly mean (the top 5% of the observed distribution)^5^. The May 2026 event doubled this threshold, making it not only a rare climatological event but one with substantial ecological significance. Climate change research has already documented trait responses to shifts in mean temperature^1,2^. Fewer studies, however, address ECEs specifically, despite their potential as major drivers of phenotypic and evolutionary change, as in the classic case of Darwin’s finches following severe Galápagos droughts^6^. Those that do, have typically focused on events that, while in the upper 5% of the observed distributions^5^, still fall within the historical climatic ranges; consequently, they may not fully capture the phenotypic or evolutionary consequences of truly unprecedented events such as the May 2026 heatwave.

To assess the consequences of this unprecedented event, we leveraged data from the Wytham Woods long-term population. We compared fledgling number and nestling mass across relative (rather than calendar) periods before and during the heatwave in 2026, against equivalent relative periods in 1978-2025 (non-heatwave years), controlling for reproductive timing itself (see Supplemental Information (SI)-1). The seasonal decline in both measures was significantly steeper in 2026 than in non-heatwave years, with no significant differences between blue and great tits (p>0.449). Fledgling number declined by 2.60±0.12 nestlings on average in non-heatwave years and by 4.48±0.88 nestlings in 2026. Nestling mass declined by 1.02±0.06 g on average in non-heatwave years and by 2.06±0.38 g in 2026 (Fig. 1C, see SI-2 for system and model details, and full results). Given the short-term nature of the heat spell, these effects most likely reflect direct impacts on parental foraging and provisioning, and on nestling thermoregulation (e.g. panting)^7^. Since fledgling mass strongly predicts overwinter survival and carries over into adult body mass^8,9^, heatwave-exposed broods produced not only fewer, but lower-quality, offspring, which may have consequences at the population level over the long term.

The 2026 heatwave coincided with the tail-end of the 2026 breeding season, but, because the May 2026 event was exceptionally severe, we expect its impacts to extend beyond late broods and have pervasive effects across the breeding season. In fact, even if early broods may have avoided the most intense effects of the heatwave in the nest, their recently fledged juveniles may have been impacted once they were no longer under parental care. The post-fledging period is a vulnerable stage for birds, with juvenile birds facing high mortality due to, for instance, limited foraging abilities^10^. A heatwave of this magnitude, which we have shown to affect nestlings under parental care, is therefore likely to further reduce the already low survival prospects of recently fledged juveniles, extending the impacts beyond active broods to much of the breeding cohort.

In conclusion, the May 2026 heatwave, unprecedented in intensity and timing, notably worsened seasonal declines in fledgling number and nestling mass, with likely longer-term effects via survival. As such extreme events increase in frequency^3^, our findings offer an early benchmark for their impacts. Continued monitoring of long-term populations remains essential for quantifying the effects of such extremely severe events and are invaluable for determining whether these acute costs we describe persist across generations.

## Acknowledgements

We thank all researchers and fieldworkers involved in the long-term monitoring of Wytham’s tit populations. This work was supported by NERC grants NE/X000184/1, NE/K006274/1 and NE/S010335/1, UKRI Frontiers grant EP/X024520/1, BBSRC grant BB/L006081/1 and ERC grant AdG250164 to BCS.

## Author contributions

DLI and BCS conceptualised the study. DLI collated, analysed the data and wrote the original draft, which was reviewed, edited and improved by DS and BCS.

## Supplemental Information (SI)

### SI 1: Distribution of breeding attempts and sample sizes per group

**SI 1-Figure 1:**
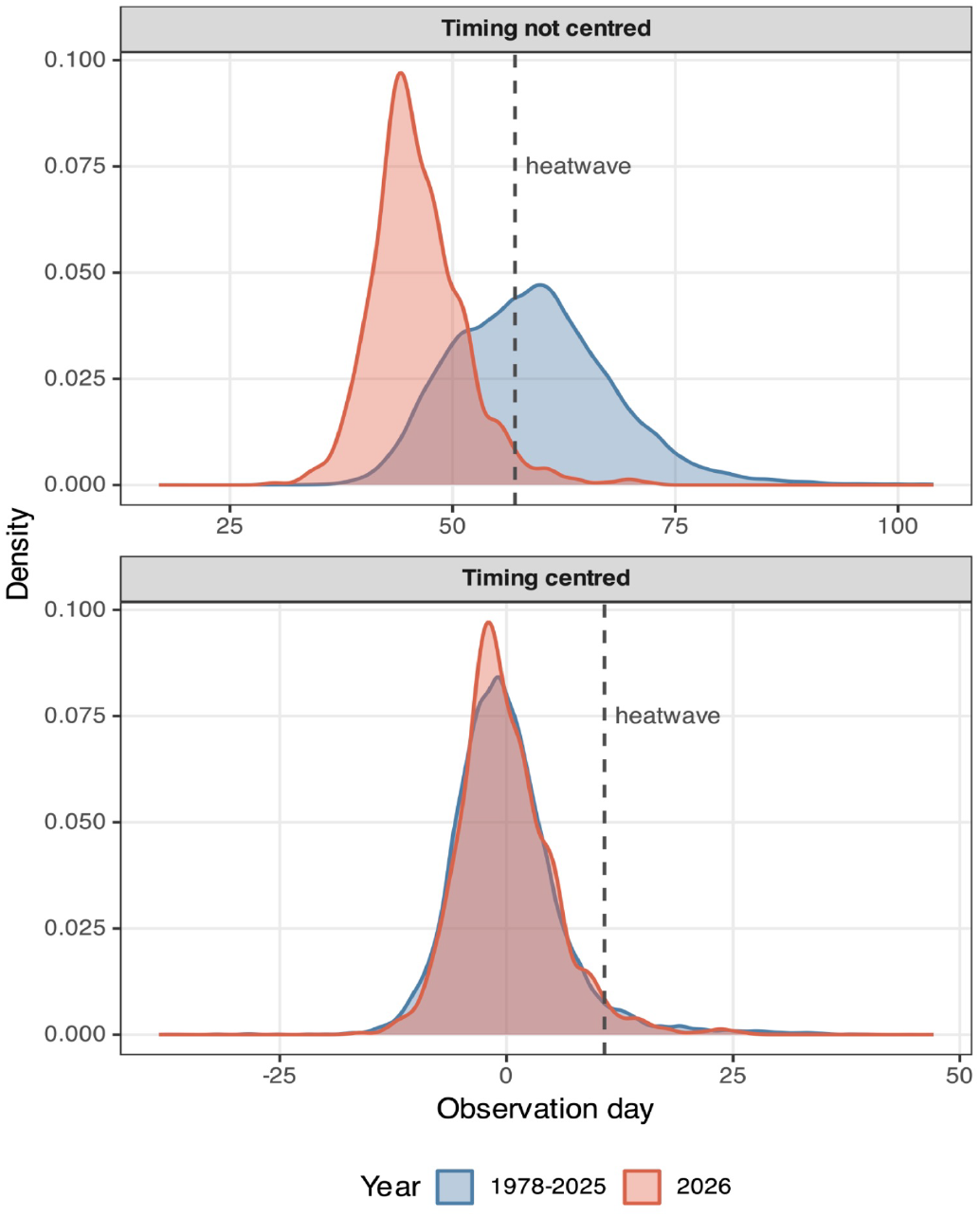
Mean centring observation day within a year allows comparison between reproductive events at the same relative time of the breeding season. In Timing not centred values in the x axis represents days since the first of April.

### SI 2: Methodological details and full model results

#### Study system

We studied the great tit (*Parus major*) and blue tit (*Cyanistes caeruleus*) population of Wytham Woods, a woodland near Oxford (51º 46 N, 1º 20’ W) where 1204 wood-crete boxes are available for the tits to breed. Wytham is divided into nine sections (hereafter: section) which, although subjectively delimited, still capture environmental differences in terms of habitat, number of boxes available and other factors that can influence great tit mass (see^1^). Between March and June, each box is visited at least once a week to collect detailed breeding and morphological data. Nestlings are ringed at 14 days old, when they are measured to record body mass (to the nearest 0.1 g^2^).

We collated hourly temperature data for the period 15-May 15-June between 2023 and 2026 from 218 temperature loggers (TOMST) placed in trees within Wytham Woods that record temperature every 15 minutes. In our analyses we used hourly mean temperature values. Furthermore, we also collated historical mean, maximum and minimum daily temperature for that period between 1772 and 2025 from the Central England temperature dataset (CET; Met Office Hadley Centre - www.metoffice.gov.uk/hadobs/). Data used to prepare Figure 1A was obtained from the HadUK-Grid 1km gridded observational dataset (v.1.3.2)^3^.

#### Statistical analyses

##### Analysing the magnitude of the heatwave

Using the thermo-logger and historical data, we compared mean hourly temperatures during the 2026 May heatwave to mean hourly values in Wytham between 2023-2025 and to mean daily values from the CET dataset between 1772-2025. We compared temperatures for two periods: the full heatwave interval from the 22nd to the 29th of May, and a shorter interval between the 25th and 27th of May, when temperature in 2026 was the highest.

##### Heatwave effects on adult reproductive success and nestling mass

We analysed the effects of the heatwave by comparing number of fledglings, a measure of reproductive success, and nestling mass in the periods before and after the 27th of May in the year with a heatwave (2026) and in years with no heatwave (1978-2025). To deal with interannual variations in reproductive timing, we defined period (pre-heat-wave vs heatwave) relative to each year’s timing rather than using absolute values. This allowed us to compare the values of the tail of the distribution in 2026 to those in a similar relative position in the breeding seasons between 1978 and 2025 (see SI 1).

We tested the heatwave effects on number of fledglings by fitting one linear mixed effects (LMM) model. The model included number of fledglings as the dependent variable and period, heatwave (heatwave year -2026-vs non-heatwave years -1978-2025-), species (great tits vs blue tits) and section as explanatory variables. We also included an interaction between period and heatwave to analyse differences in breeding success after experiencing a heatwave, and a triple interaction between period, heatwave, and species to test for species-specific responses. As random effects we included nestbox, mother identity, and year.

To test the effects of the heatwave on nestling mass we fitted an additional LMM that included nestling mass as a dependent variable. As fixed effects the model included period, heatwave year, species, section and clutch size. In addition, we also included a triple interaction between period, heatwave year and species. In this model, as random effects, we included mother, brood identity, and year.

All models were fitted in R v. 4.5.2^4^ using the packages *lme4* v. 1.1-37^5^ and *lmerTest* v. 3.1-3^6^. Model residuals were visually inspected and showed no marked deviations from normality. Non-significant triple interactions (p>0.05) were removed from the models. Pairwise comparisons between factor levels were conducted using Tukey HSD correction with the package *emmeans* v. 2.0.0^7^

**SI 2 Table 1.**
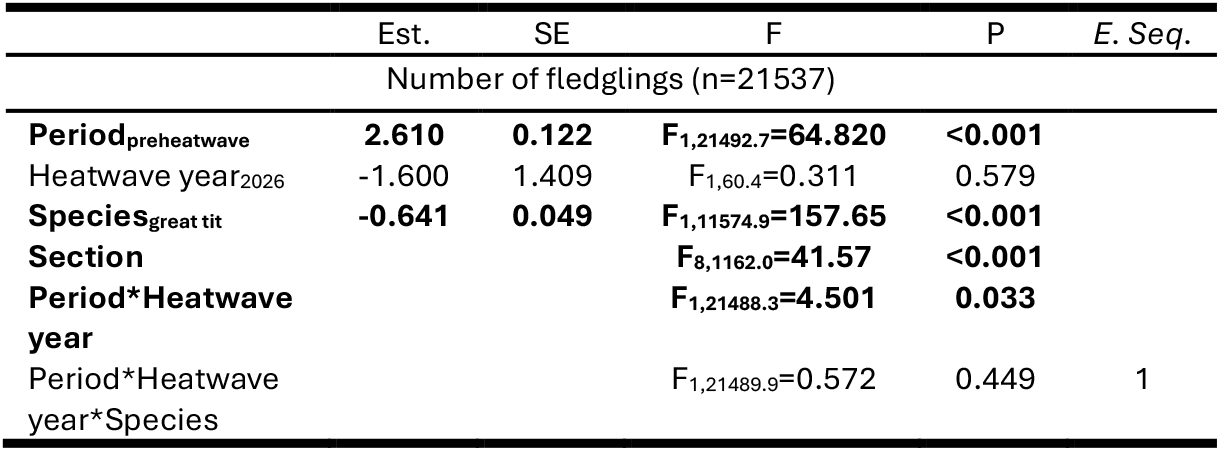
Results from the linear mixed effect model analysing the effects of the heatwave on number of fledglings (mean centred). Bold denotes statistical significance. Reported values from excluded variables refer to the step before their exclusion.

**SI 2 Table 2.**
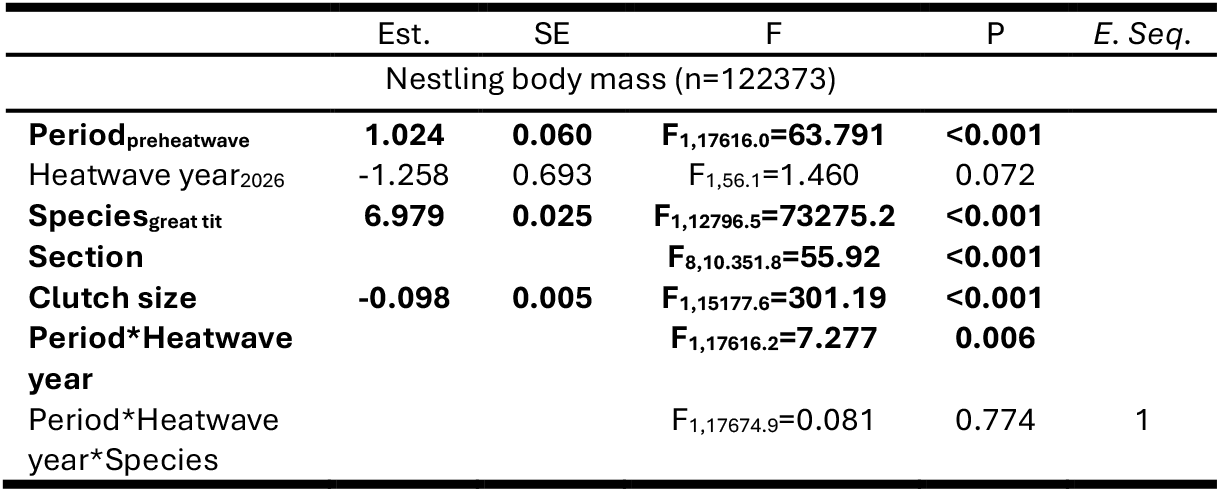
Results from the linear mixed effect model analysing the effects of the heatwave on nestling mass (mean centred). Bold denotes statistical significance. Reported values from excluded variables refer to the step before their exclusion.

